# LncRNAs of *Saccharomyces cerevisiae* dodge the cell cycle arrest imposed by the ethanol stress

**DOI:** 10.1101/2021.06.28.450142

**Authors:** Lucas C. Lázari, Ivan R. Wolf, Amanda Piveta Schnepper, Guilherme T. Valente

## Abstract

Ethanol impairs many subsystems of *Saccharomyces cerevisiae*, including the cell cycle. Cyclins and damage checkpoints drive the cell cycle. Two ethanol-responsive lncRNAs in yeast interact with cell cycle proteins, and here we investigated the role of these RNAs on the ethanol-stressed cell cycle. Our network dynamic modeling showed that the higher and lower ethanol tolerant strains undergo a cell cycle arrest during the ethanol stress. However, lower tolerant phenotype arrest in a later phase leading to its faster population rebound after the stress relief. Two lncRNAs can skip the arrests mentioned. The *in silico* overexpression of lnc9136 of SEY6210 (a lower tolerant strain), and CRISPR-Cas9 partial deletions of this lncRNA, evidenced that the one induces a regular cell cycle even under ethanol stress; this lncRNA binds to Gin4 and Hsl1, driving the Swe1p, Clb1/2, and cell cycle. Moreover, the lnc10883 of BY4742 (a higher tolerant strain) interacts to the Mec1p and represses Bub1p, circumventing the DNA and spindle damage checkpoints keeping a normal cell cycle even under DNA damage. Overall, we present the first evidence of the direct roles of lncRNAs on cell cycle proteins, the dynamics of this system in different ethanol tolerant phenotypes, and a new cell cycle model.

## INTRODUCTION

Worldwide ethanol production relies on the use of *Saccharomyces cerevisiae* (Gupta and Verma 2015; Mohd Azhar et al. 2017). However, the high ethanol yield challenges the production stalling the fermentation (Kasavi et al. 2016). Ethanol stress alters many yeast pathways dampening the yeast viability, cell growth, macromolecules biosynthesis, fermentation, and membrane integrity (Ohta et al. 2016; Auesukaree 2017). The yeast ethanol-stress responsive mechanisms and the cell surveillance comprise the regulation of plenty of specific genes influencing a wide range of biological processes, especially the cell cycle and viability pathways (Chandler et al. 2004; Teixeira et al. 2009; Li et al. 2017). For instance, the activity of several cell cycle-related genes (e.g., BUB1, CDH1, CLN3, SWE1, GRR1, SWI4, and SWI6) is crucial to keep the yeast cell growth in a medium with 11% of ethanol (Kubota et al. 2004). Altogether, knowledge concerning the dynamic of cell cycle systems in stressful conditions is essential to develop new ethanol tolerant strains.

Cyclins activate kinases to guide the whole cell cycle. The G1 phase activates the transcription factors causing the cell cycle *START*, in which cells reach a constant growth and protein synthesis leading to the activation of Cln1p, Cln2p, or Cln3p cyclins. Activated cyclins bind to the Cdc28p kinase activating the transcriptional factors SBF and MBF, which control the transcription of essential genes for subsequent phases (Bloom and Cross 2007). The genome duplicates at the S phase via Clb5p and Clb6p type B cyclins activation. Additionally, conjoint action of Clbs and pre-replicative complex are essential for proper DNA unpacking and polymerases recruitment (Nougarede et al. 2000; Tanaka and Diffley 2002; Sheu and Stillman 2010); then, cells may be ready to shift to G2. After duplication of spindle pole bodies by the joint action of Clbs proteins (Clb1p, Clb2p, Clb3p, and Clb4p), the activation of cyclin inhibitors is crucial to degrade Clbs, prompting the anaphase completion (Bloom and Cross 2007). Throughout the phases, checkpoint mechanisms halt the cycle progression in damaged cells until anomalies repair, e.g., inappropriate cell size, DNA damage, unattached kinetochores, misaligned spindle, and mating (Wang et al. 2000; Hu et al. 2001; Yasutis and Kozminski 2013; Barnum and O’Connell 2014).

Long non-coding RNAs are ncRNAs >200 nucleotides length that works on many regulatory processes (Lee 2012; Yamashita et al. 2016; Engreitz et al. 2016; Han et al. 2017; Peng et al. 2017). LncRNAs can interact with proteins closer to transcription sites regulating the gene expression, scaffold protein-complex, interact with chromatin complexes acting on enhancers (Ferrè et al. 2016). We previously defined the highest ethanol tolerance level for six yeast strains, and we found their ethanol-stress responsive lncRNAs. Predictions of lncRNA-protein interaction evidenced that these lncRNAs work on many strain-specific responsive systems, including the cell cycle. Among these lncRNAs, we found that the lncRNAs lnc9136 and lnc10883 of SEY6210 and BY4742 strains, respectively, bind to cell cycle-related proteins (Marques et al. 2021). Moreover, we identified many cell cycle-related genes differentially regulated during the severe ethanol stress (unpublished data).

Most of the knowledge concerning the eukaryote cell cycle is based on yeast systems. However, the dynamic of the yeast cell cycle under the ethanol stress, the role of lncRNAs along the cycle or after the stress relief, and lncRNA-cell cycle protein interactions are unknown. Herein we addressed these points using dynamic network modeling, population rebound experiments, and CRISPR-Cas9 approaches. We developed a logic model of the yeast cell cycle to assess the ethanol stress influence and the molecular basis of lnc9136 and lnc10883 in the cell cycle progression. We used transcriptome data of lower (LT) and higher (HT) ethanol tolerant strains under severe ethanol stress to perform the dynamic network modeling. The modeling showed that both phenotypes undergo a cell cycle arrest during the extreme ethanol stress, albeit two lncRNAs can circumvent these arrests. LTs arrests in a later stage of the cell cycle (M phase) than HTs (G1 phase), which may be responsible for the faster population rebound after the ethanol stress relief observed for the earlier phenotype in our rebound experiments. Remarkably, we associated a positive impact of lnc9136 of SEY6210 (an LT strain) in its cell cycle. In this case, overexpressing this lncRNA rescued the expected arrest by repressing two proteins, allowing SEY6210 to head away in the cell cycle, even under ethanol stress; the partial deletion of this lncRNA by CRISPR-Cas9 corroborates this finding. Finally, we associated the lnc10883 of BY4742 (an HT strain) to the lack of DNA and spindle damage checkpoint, keeping the cell cycle, even though cells harbor DNA damage.

## MATERIAL AND METHODS

### Network designing

KEGG cell cycle pathway (sce04111) was the scaffold network of our logic model. The nodes and edges represent variables (e.g., proteins, genes, and events) and interactions (activation or inhibition) structured using the GINsim (Chaoyia et al. 2011). Nodes’ value changes during the simulation based on the node-regulator interactions described in the logical functions (further described). Missing proteins and interactions were added based on literature data. We used phenomenological nodes (MASS, BUD, DNA_REPLICATION, SPINDLE, and MITOSIS_EXIT) to track the cell cycle progression and model cyclic behavior. These nodes indicate whether the mass, budding process, DNA replication, and spindle assembling are adequate for cell division and whether the model launched the mitosis (Fauré et al. 2009) (**Figure 1**).

**Figure 1:**
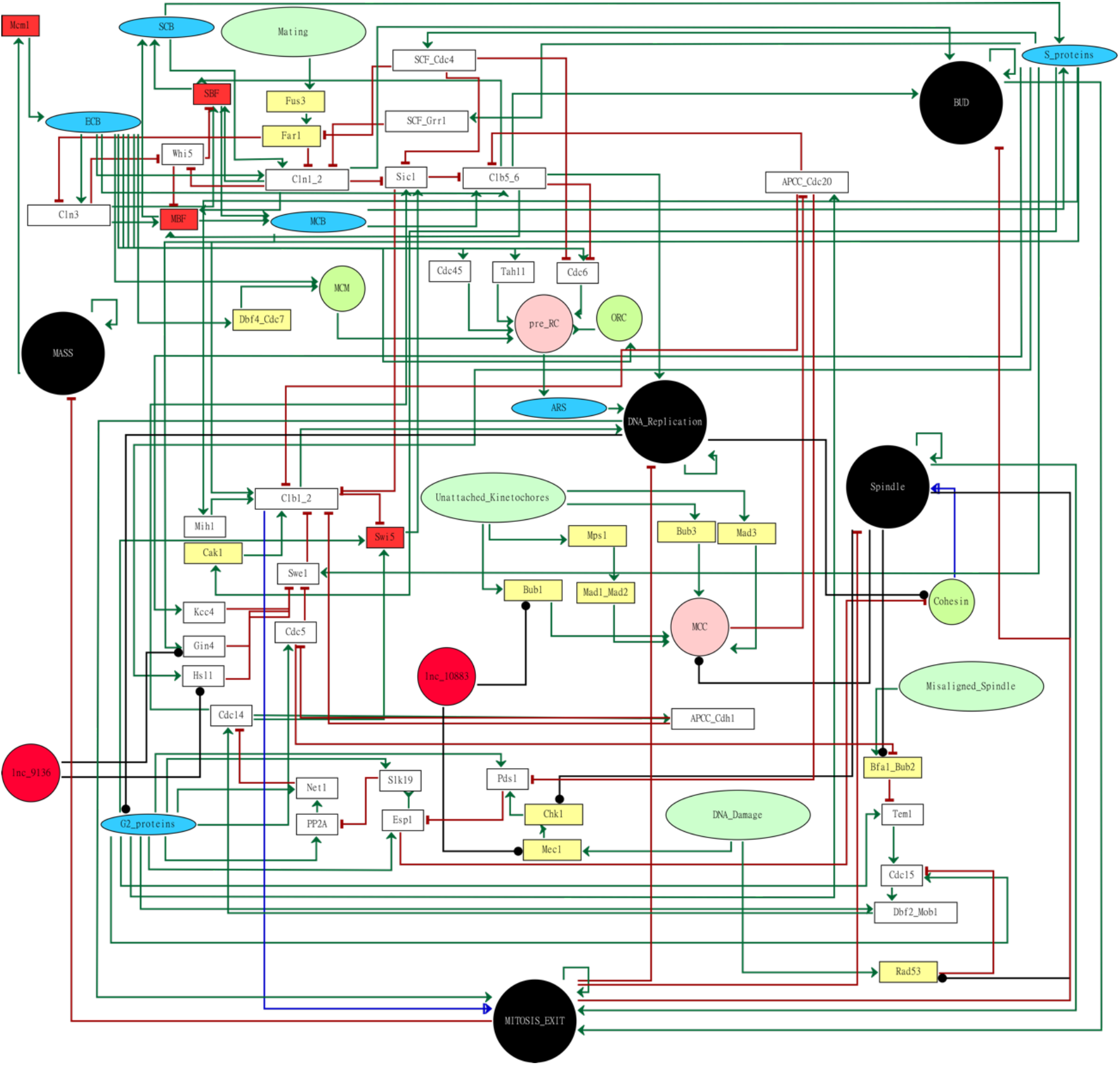
Network model. **Node colors**: purple are protein complexes, black are phenomenological nodes, red are transcription factors, green ellipses are checkpoints, white are single proteins, and green circles are protein complexes. The whole computational GINsim model is available in https://figshare.com/account/projects/112128/articles/14503035.

Protein complex formation was implicitly assumed since logical modeling is not able to deal with mass flow. Thereby, the complex activation relies on all components’ presence; complexes’ level changes during the simulation according to the smallest value in the component. Complexes with subunits without regulatory changes through the cell cycle are not in the networks, such as previously performed (Fauré et al. 2009).

We created logical equations for each interaction (further described) to drive the value shifts of regulated targets (**Supplementary Table 1-2**). The model was adjusted to simulate the normal cell cycle behavior: mass increasing, the cell cycle START activation, progression through phases, mitosis exit, and mass decreasing. The model performance was assessed by simulating 109 cell cycle mutant phenotypes (knockouts and overexpression) published (**Supplementary Table 3**). Therefore, we added these absent interactions in the model, followed by adjustments of logic functions, to fit the mutant phenotypes. The model robustness was checked by simulating the cell cycle from 100 random inputs (initial node values).

### Development of logical functions

Cell cycle progression throughout the phenomenological nodes, their activation, and checkpoints were modeled in GINsim (Chaoyia et al. 2011) using logical functions with Boolean operators AND, NOT, and OR (**Supplementary Table 1; https://figshare.com/account/projects/112128/articles/14503035**).

Each node in the model can have 1 out of 4 values, which “0”, “1”, “2”, and “3” indicates absence, low, normal, and high quantities, respectively; for phosphorylated or de-phosphorylated proteins, the values designate the amount of active forms. Nodes poorly explored in the literature (e.g., CAK1, MIH1, and mating nodes) were limited to “0” or “1” to avoid spurious logical functions. Nodes corresponding to the down-regulated genes (further described) were also limited to “0” or “1”, indicating the absence or low protein quantity, respectively. Conversely, we enhanced the level of activators of up-regulated genes to a certain level to allow regulation (inhibition or higher activation) by other nodes (**Supplementary Table 1-2**).

The values associated with phenomenological nodes indicate activity rather than abundance. For instance, the “BUD” node may have “0”, “1”, or “2” denoting no budding, activation, or formation, respectively. Phenomenological nodes keep the maximum value until suppression by the MITOSIS_EXIT node.

Based on the literature data, the activators and their regulated node levels are positively or negatively related to activation or inhibition, respectively. Moreover, an inhibitor must have a higher value than an activator to inhibit the regulated node and vice versa. Protein phosphorylation was also modeled as mentioned.

### Model cycling rationale

Dynamic logical models reflect state transitions rather than quantities (Fauré et al. 2009). The mass strongly influences cell cycle START. Conversely to other nodes, the node MASS is self-activated in the model (MASS="1"). The maximum MASS value triggers the cell cycle START by activating the MCM1 (Mcm1p regulates the transcription of G1 and S genes (Chang et al. 2017)); all other nodes’ values start with “0".

The cell cycle modeling starts activating Clns inducing SBF, MBF, and BUD (bud formation starts) nodes activation. Then, the model activates the pre-replication complex triggering the DNA replication (DNA_REPLICATION node). This state persists active until the end of mitosis. We set the DNA_REPLICATION node as a G2 to M transition triggers by activating the G2_proteins node. The G2_proteins node keeps the progression tracking and starts several nodes related to the M phase at the right time. The lack of “0” level in the MASS node after the beginning of simulation denote the cell cycle arrest (e.g., the “X” axes of **Figure 3–5; Supplementary Figure 1**).

The activation of the MITOSIS_EXIT node (value=“1”) starts the M phase and requires activation of all phenomenological nodes (except MASS) and Clb1/2. The anaphase and mitosis exit rely on the complete Clb1/2 inhibition. Hence, the MITOSIS_EXIT is “2” and inhibits all phenomenological nodes, restarting the cycle.

Conversely to other nodes, the checkpoint ones (DNA_DAMAGE, MISALIGNED_SPINDLE, UNATTECHED_KINETOCHORES, and MATING) were the model’s inputs to allow checkpoint outcomes. Slight and strong activation/response of those checkpoints (except the MATING) were designated as “1” and “2”, respectively. Checkpoint reaching “2” forces the cell cycle arrest by inhibiting crucial nodes for cell cycle progression (**Figure 1**). Negative feedback from the SPINDLE node may deactivate checkpoint nodes below “2". Since the matting checkpoint is needless for us, we set the MATTING node as Boolean: its activation arrests at G1.

### Simulating the impact of ethanol stress in cell cycle

There is a high correlation between RNA and protein levels in yeasts (Skelly et al. 2013); thereby, we assumed the protein levels in our simulations based on the transcriptome data. We previously tested the highest ethanol tolerance level of six *S. cerevisiae* strains. Therefore, three strains fit a higher tolerant (HT) and three a lower tolerant (LT) phenotypes (Marques et al. 2021). The Log2 fold-change comparing treatment (the highest ethanol level supported for each strain) vs. control were retrieved from previous RNA-Seq data (unpublished data) (NCBI BioProject number PRJNA727478). These expressions of cell cycle network genes were used to simulate the impact of the severe ethanol stress on HTs’ and LTs’ cell cycles. In this case, we selected transcripts with similar expression profiles (up or down-regulation) in all three strains within each phenotype. We also performed simulations to compare the impact of ethanol on DNA damage checkpoint amongst strains using expression data of this pathway (**Supplementary Table 2, and 4**).

LncRNAs may activate complex formation or inhibits target proteins, acting as scaffolders or as protein baits, respectively (Chen et al. 2019). To assess the impact of ethanol stress-responsive lncRNAs in the cell cycle, we embedded the lncRNA lnc9136 of SEY6210 (an LT strain) and lnc10883 of BY4742 (an HT strain) into the network (**Figure 1**).

The lnc9136 interacts with Gin4p and Hsl1p, while the lnc10883 interacts with Mec1p and Bub1p (**Figure 1**); lncRNAs’ genomic coordinates and sequences are in the **Supplementary Data 1** and NCBI accession numbers MZ099632, and MZ099633. First, based on transcriptome data, we tested the logical functions stating the lncRNAs mentioned either as activator or inhibitor due to the lack of information about their effects on target proteins. Hence, we assessed the lncRNAs impact on the cell cycle by increasing or decreasing the target protein levels without adding excessive model uncertainty. Since the lnc10883 interacts with checkpoint proteins, we *in silico* overexpressed this lncRNA to appraise its impact on the spindle damage pathway under the hypothesis that stressed strains are undergoing spindle damage (**Supplementary Table 2**).

### Partial deletion of lncRNA 9136

To assess the modeled hypothesis regarding the positive influence of lncRNA lnc9136 of SEY6210 under the severe ethanol stress (further discussed), we generated partial deletion mutants (hereafter referred to as lnc9136Δ1, and lnc9136Δ2) of this gene using CRISPR-Cas9 (**Table 1; Supplementary Figure 2**). The pMEL16 (His-, Addgene 107922) and p414-TEF1p-Cas9-CYC1t (hereafter referred to as P414, Addgene 43802) were the plasmids used to express the gRNA and Cas9, respectively. The P414 has the KAN selective marker rather than TRP1 (donation from Dr. Arnold Driessen of the University of Groningen, The Netherlands).

**Table 1:**
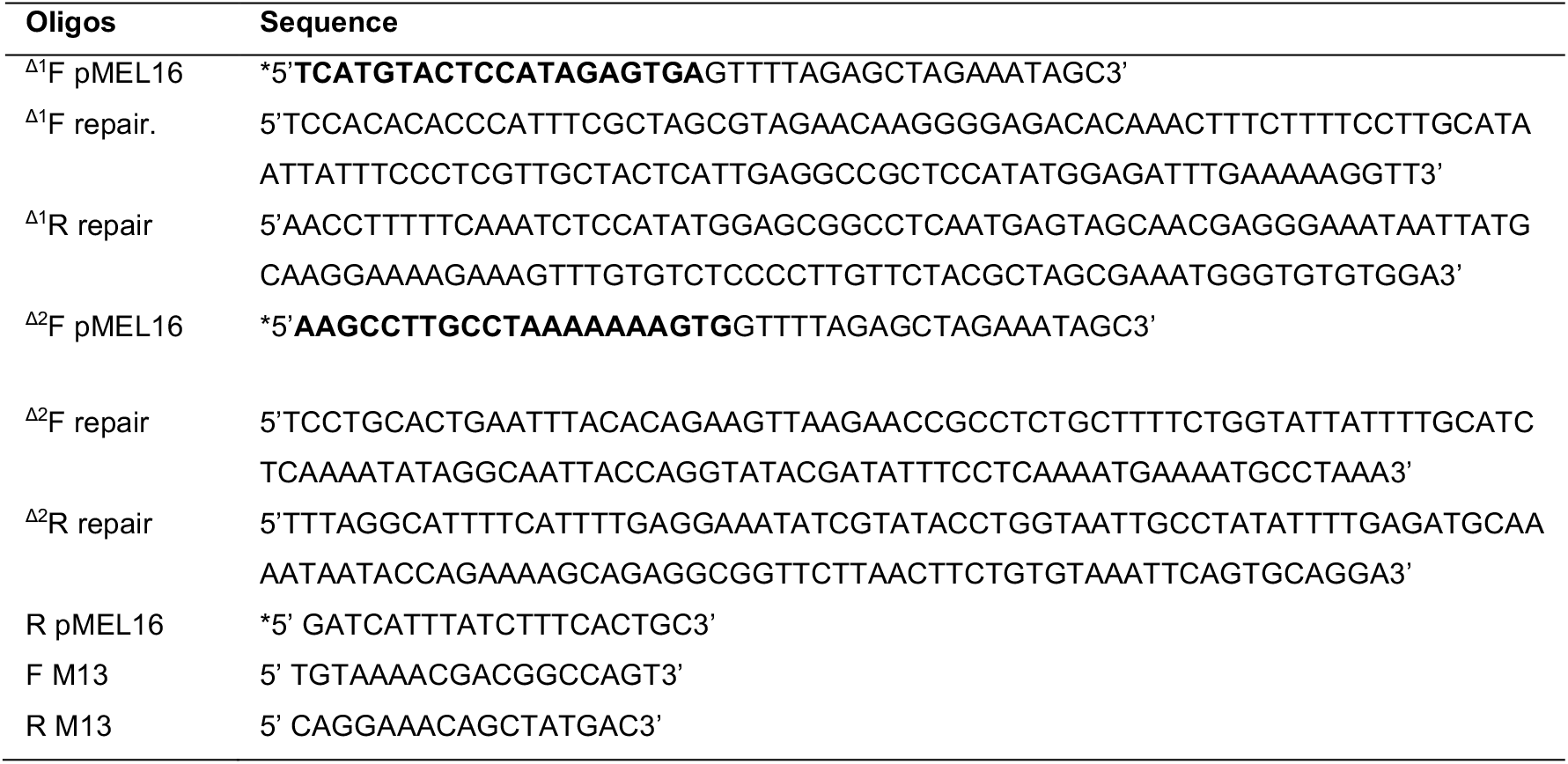
Oligos used in this work. Bold nucleotides are related to the genomic target sequence. **“Δ1”**: oligos to obtain the SEY6210 lnc9136Δ1; **“Δ2”**: oligos to obtain the SEY6210 lnc9136Δ2; *: phosphorylated ends. Details of target-homology repairs and Cas9 cleavage loci are in **Supplementary Figure 2**.

The target region was inserted into pMEL16 by PCR: 20 ng pMEL16 plasmids, 1 μL of Phusion (NEB M0530S), 1X of Phusion Buffer, 0.1 mM of each dNTP, 0.4 μM of F pMEL16 and, 0.4 μM of R pMEL16 in a final volume of 25 μL (**Table 1**). Touchdown reaction was 98°C for 1 min, followed by 5 cycles with a higher Tm (98°C for 30 sec, X°C for 30 sec, and 72°C for 6 min), 10 cycles with medium Tm (98°C for 30 sec, Y°C for 30 sec, and 72°C for 6 min), 20 cycles with lower Tm (98°C for 30 sec, Z°C for 30 sec, and 72°C for 6 min), and 72°C for 6 min. The X°C, Y°C and Z°C annealing temperatures are the averages between the melting temperature of R pMEL16 and the F pMEL16 of each gene, plus 9°C, 5°C and 2°C, respectively. Then, 25 μL of amplicons were digested with 1 μL of DpnI (NEB R0176S). 100-200 ng of purified digestion was ligated by T4 DNA ligase (Promega M1801), and 2 μL of the product were transferred into 40 μL of TOPO competent cells (ice incubation for 20 min, 42°C for 50 sec, and heat shock on ice for 2 min). Cells were incubated in 200 μL of LB medium at 37°C for 1h, followed by overnight incubation (37°C) on an LB plate with 0.05 mg/mL of ampicillin. Colonies with modified pMEL16 were sought by PCR-RFLP using M13 oligos (**Table 1**) and ClaI digestion (Bsu16l, Thermo Fisher IVGN0306) (37°C for 1h); positive colonies were grown in liquid LB with 0.05 mg/mL of ampicillin, and plasmids were extracted using the QuickLyse Miniprep system (Qiagen 27405).

Yeast is overnight grown (30°C) in liquid YPD (peptone 2%, yeast extract 1%, and glucose 2%), diluted in the same medium to an OD600 of 0.3, and incubated (30°C, and 200 RPM) until OD600 of 1.0. Competent cells were obtained using the Yeast Transformation Kit (Sigma YEAST-1KT).

A solution with 10 μL of salmon testes DNA, 600 μL of plate buffer (both from Sigma YEAST-1KT kit), 1 μg of P414 plasmid, 1 μg of modified and purified pMEL16, 5 μL of double-strand repair DNA (**Table 1**), and 100 μL of competent yeast cells were incubated at 30°C for 30 min, and 10% of DMSO was further added. The samples were immediately incubated at 42°C for 15 min, quickly transferred into ice, and chilled for 2 min. The double-strand repair DNA was obtained by mixing 100 μM of F and R repair oligos for each target loci (**Table 1**), followed by incubation (95°C for 10 min) and slow cooling on a bench.

Cells were harvested by centrifugation (2000 RPM for 30) sec, and the supernatant was discarded by careful pipetting. The pellet was diluted using 250 μL of drop-out medium His^−^ (Yeast Synthetic Drop-out Medium Supplement without Histidine, Sigma Y1751) at 1.92 mg/mL initial concentration, supplemented with 20% of glucose, and 1.9 mg/mL of Yeast nitrogen base without amino acids and ammonium sulfate. Tubes were incubated (30°C, 200 RPM for 2h), followed by plating on drop-out medium His^−^ (the same one mentioned, plus 2% of bacto agar, and 0.2 mg/mL of G418). Plates were incubated at 30°C until colonies arise. Mutants were screened by standard colony PCR and sequenced by the Sanger method. Mutants were grown in liquid YPD with 0.2 mg/mL of G418. Finally, 700 μL of mutant cells plus 15% of glycerol were stored at −80°C until use.

### Population rebound experiments

We investigated the population rebound of SEY6210 lnc9136Δ1 mutant and SEY6210 wild-type (WT) after the high ethanol stress relief. Overnight cells grown in YPD at 30°C (YPD plus 0.2 mg/mL of G418 for mutants) were diluted in YPD to an OD600 of 0.3 and incubated (30°C, and 200 RPM). Then, OD600 0.2 cells (~400 μL of cells) were harvested by centrifugation (2000 RPM for 2 min), and the medium was carefully discarded by pipetting. Pellet cells were diluted in 2 mL of pre-prepared YPD with different ethanol concentration (18%, 20%, 22%, 24%, and 26% (v/v)). The content was transferred into 10 mL rounded bottom tubes and immediately incubated (30°C, 135 RPM for 1h). Then, 1 mL of cells were harvested by centrifugation (2000 RPM for 2 min), the medium was carefully discarded by pipetting, and the pellet was diluted in 1 mL of YPD. Finally, 200 μL of each sample were transferred into each well plate in four technical replicates. Plates were incubated at 30°C, and OD600 was measured each five min along 24h before 30 sec of orbital shaking. The same experiment was also performed to compare the BY4742 and SEY6210 wild-types under 26% and 20% of EtOH stress (their highest ethanol tolerance levels (Marques et al. 2021)). We compared the Log *k* between SEY6210 WT vs. SEY6210 lnc9136Δ1 mutant and between SEY6210 vs. BY4742 wild-types using the mixed-effects model (Geisser-Greenhouse correction, Sídák test, and swap direct comparisons), and the Mann-Whitney U (unpaired mode, non-parametric, and two-tailed) tests, respectively.

## RESULTS

### The model’s performance

The network has 67 nodes and 144 interactions (**Figure 1**). A total of 82 out of 100 random inputs simulations outcome a cyclic attractor passing throughout all cell cycle phases. Thus, the largest basin of attraction (composed of these 82 random inputs) is a functional cell cycle. The logical functions of the model are available in **Supplementary Table 1-2**.

The model reaches 86.6% accuracy simulating the data of cell cycle imbalance mutants. Some papers about these mutants did not mention the cell cycle phase arrested impairing our confusion matrix: e.g., many simulations gave out an arrest, albeit the respective mutants were previously described simply as “Inviable”. We considered these cases as correctly predicted by our model. We settled them as “Inviable” to calculate the accuracy, albeit it shrank the sensitivity of some classes. However, all simulated classes presented good sensitivity and specificity, being the “Arrest in G1” the only exception due to the mentioned reasons (**Figure 2A; Supplementary Table 3**).

**Figure 2:**
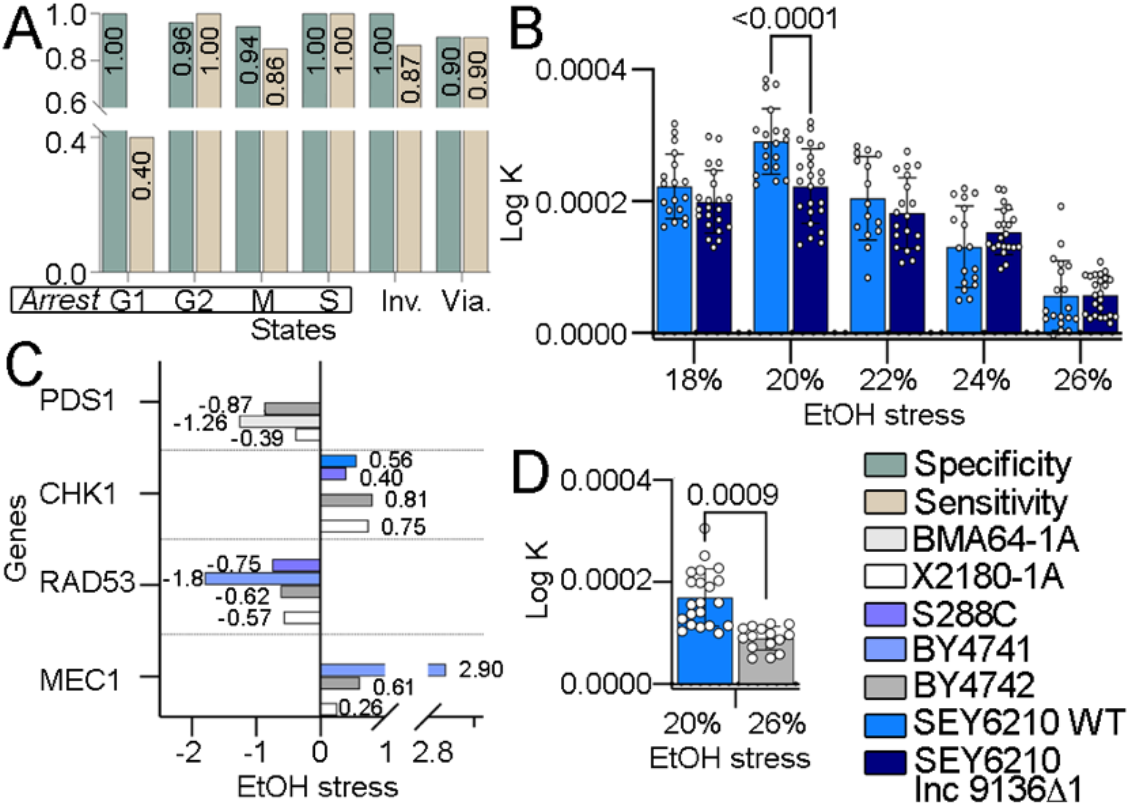
Model evaluation, growth curves, and gene expression data. **A:** simulations performance on mutants data. “Inv.” and “Via” mean “Inviable” and “Viable” phenotypes, respectively; **B:** growth curve analysis of SEY6210 WT and SEY6210 lnc9136Δ1 in the population rebound experiment after the extreme ethanol stress; **C:** Log2 fold-change of genes related to DNA damage pathways comparing treatment vs. control under the highest ethanol stress level for each strain; **D:** growth curve analysis of SEY6210 and BY4742 wild-types in the rebound experiment after the highest ethanol stress level for each strain (“X” axis); The dots at “B” and “D” are either technical or biological replicates.

### Simulating the impact of ethanol stress on cell cycle

The simulations using the transcriptome data of yeast under the highest ethanol stress (**Supplementary Table 2**) showed that HT and LT phenotypes undergo a cell cycle arrest during this stress. The HTs arrest at G1 due to the high activity of the SCF complex (its overexpression leads to an inviable phenotype), whereas the LTs arrest at the M phase (likely at anaphase) by the lack of Clb1/2p degradation/inhibition (**Figure 3**). The experiments of population rebound after the highest ethanol stress relief showed a significantly better rebound for the LT SEY6210 WT compared to the HT BY4742 WT (**Figure 2D**).

**Figure 3:**
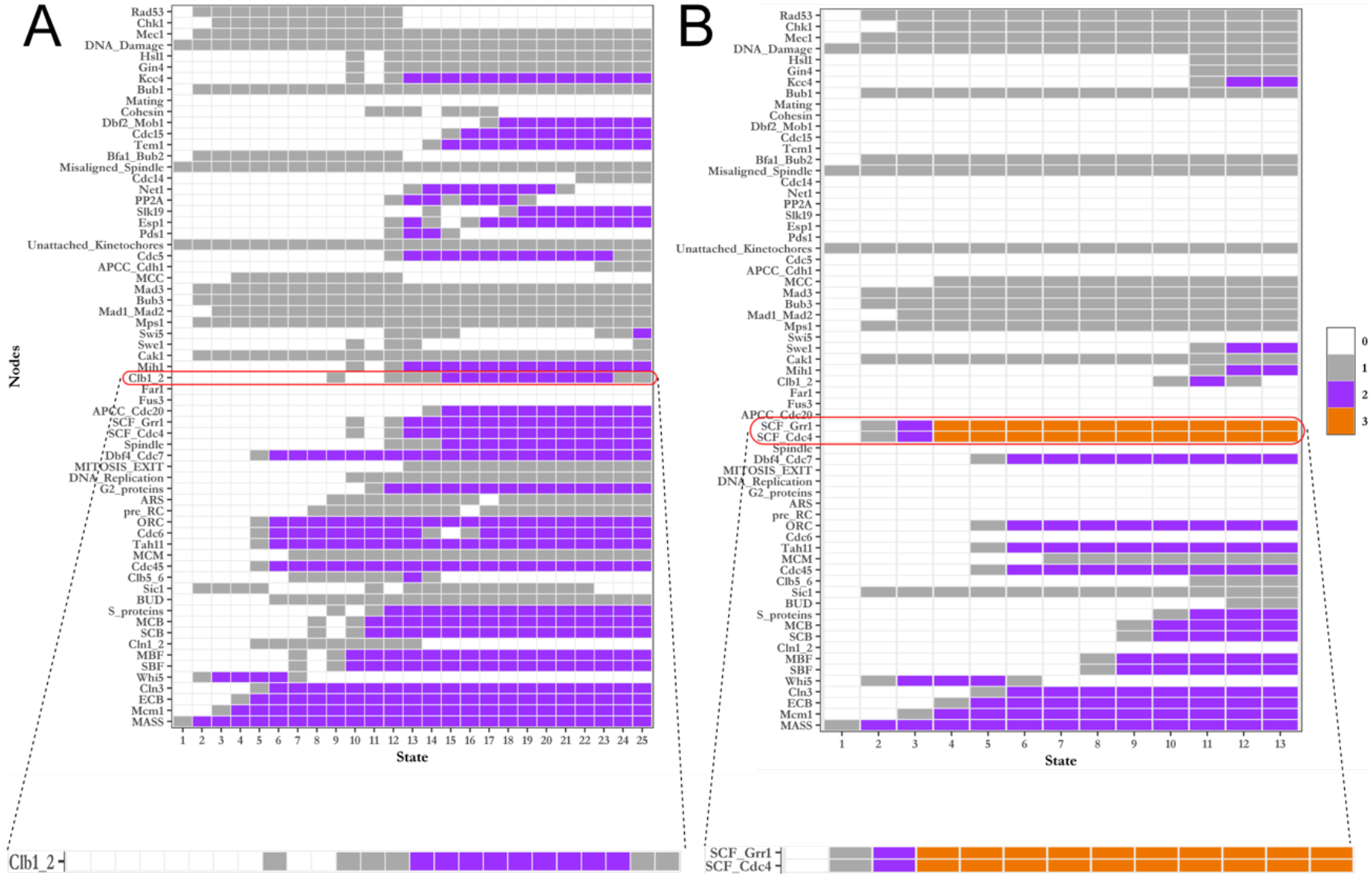
Simulation of LTs (A) and HTs (B) cell cycle under ethanol stress. The box colors indicate different levels of each node along the cell cycle. The cell cycle arrest is defined when the mass node does not return to zero. The magnified boxes indicate the nodes responsible for the cell cycle arrests.

Only simulations defining lncRNAs as target-inhibitor changed the profile of ethanol-stressed cell cycles; indeed, lncRNAs usually work as inhibitors.

The *in silico* overexpression of lnc9136 of SEY6210 (LT) at G2 skips the M phase arrest (this arrest is an LT feature). This overexpression inhibits the Hsl1p and Gin4p (which interacts with lnc9136), driving the Swe1p release to complete the cell cycle (**Figure 4**). The rebound of SEY6210 lnc9136Δ1 mutant (**Supplementary Figure 2**) is worse than WT after most ethanol stress-tested, albeit only the differences at 20% of ethanol (v/v) are statistically significant; the mutant grown better than WT at 24%, and both genotypes had similar growth at 26% of ethanol (**Figure 2B**). Notably, the SEY6210 lnc9136Δ2 mutant is inviable (**Supplementary Figure 2-3**).

**Figure 4:**
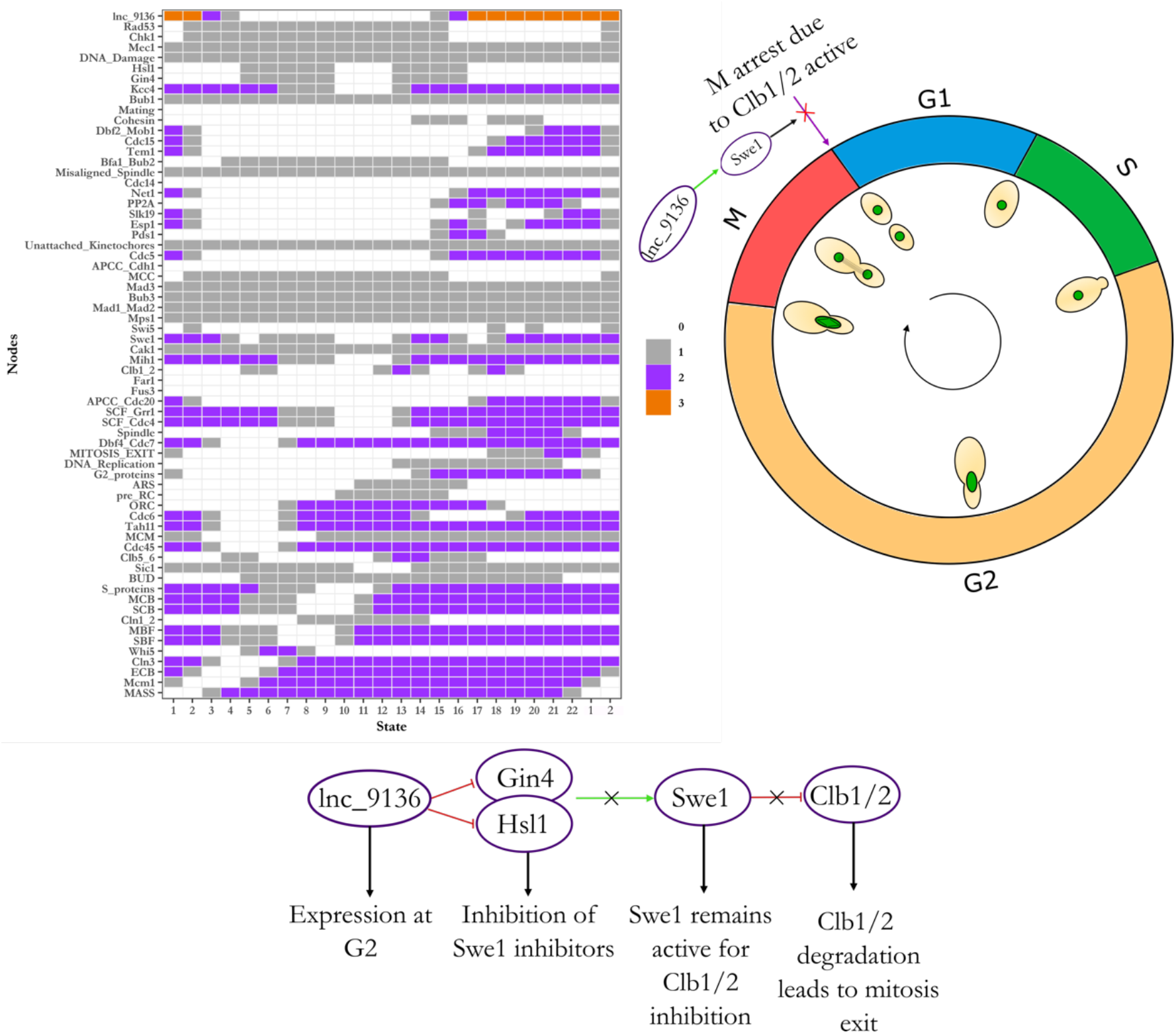
Simulation of SEY6210 cell cycle arrest skipping by overexpressing the lnc9136 at G2. The box colors indicate different levels of each node along the cell cycle. The cell cycle arrest is defined when the mass node does not return to zero. Only states “1” and “2” were plotted on the left, albeit the simulation keeps cycling through all states. The scheme on the right side depicts how the lnc9136 leads to Clb1/2 degradation. The red “X” depicts the arrest skipping.

Although the *in silico* overexpression of lnc10883 of BY4742 does not skip the G1 arrest (this arrest is an HT feature), in this case, this lncRNA inhibits its target (the Bub1p), keeping the MCC (mitotic checkpoint complex) inactive. Therefore, the APC/C complex (APCC node in our model) activates the FEAR pathway leading to the Clb1/2p degradation at anaphase, even likely with a damaged spindle. The inhibition of the Clb1/2 node enables the MITOSIS_EXIT node to reach “2", pushing the model to the end of mitosis and resetting all phenomenological nodes (**Figure 5**).

**Figure 5:**
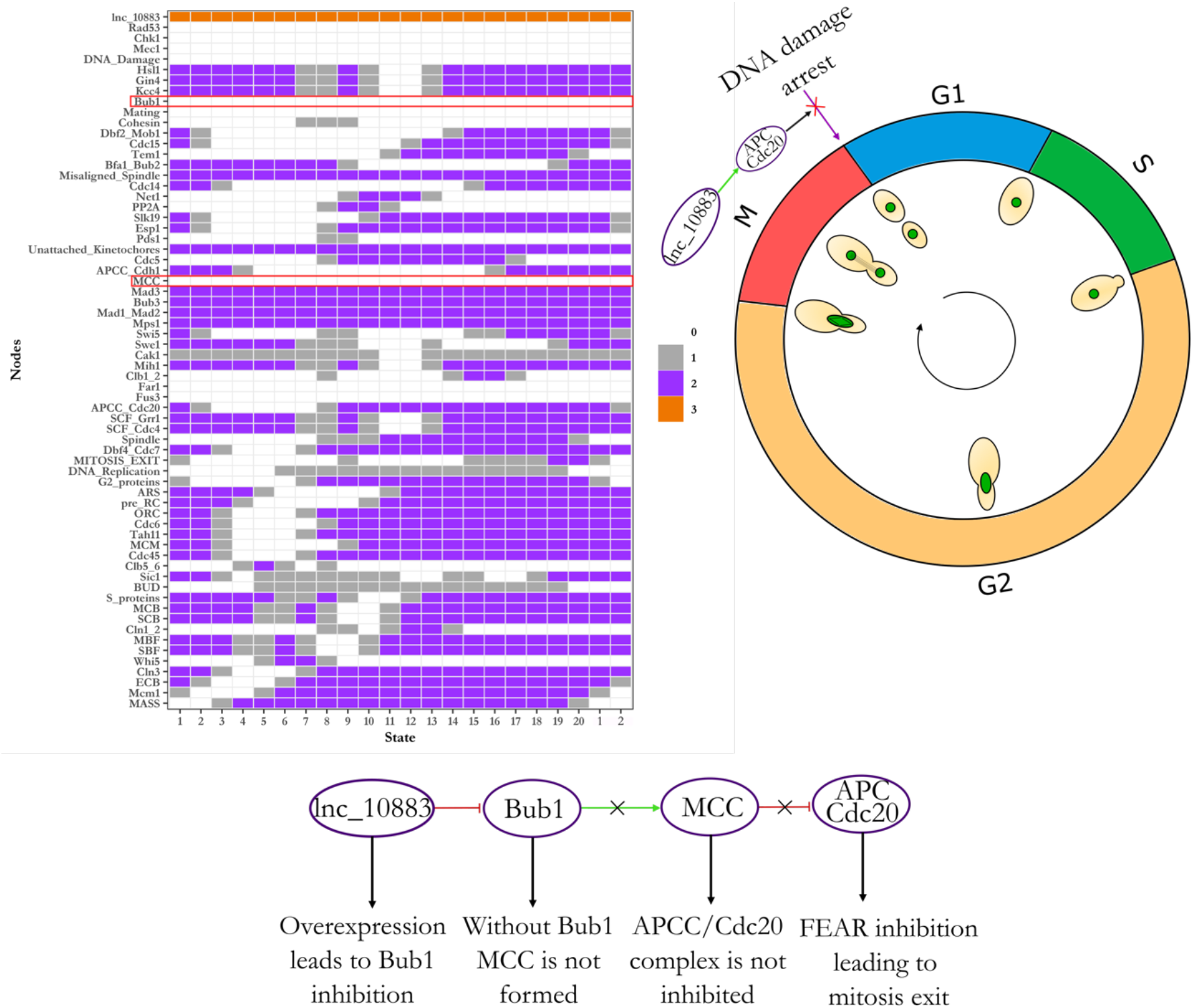
Simulation of BY4742 cell cycle overexpressing the lnc10883 and inhibiting Bub1p. The box colors indicate different levels of each node along the cell cycle. The cell cycle arrest is defined when the mass node does not return to zero. Only states “1” and “2” were plotted on the left, albeit the simulation keeps cycling through all states.The scheme on the right side depicts how the lncRNA leads to inhibiting the spindle damage response. The red “X” depicts an arrest skipping.

Previously, we found that the severe ethanol stress induces the DNA damage and the differential expression of DNA damage checkpoint-related genes in all strains (unpublished data) (**Figure 2C**). Simulation using those expression data showed that all LTs (S288C, BY4741, and SEY6210) had a cell cycle arrest, indicating a functional checkpoint. Conversely, HTs (X2180-1A and BY4742) do not stop the cell cycle progression to repair the damaged DNA (**Supplementary Figure 1; Supplementary Table 2**). The BMA64-1A was not simulated because it did not present MEC1, CHK1, or RAD53 differentially expressed.

The simulations showed that lnc10883 of BY4742 also acts on the DNA damage checkpoint by interacting with Mec1p (this protein activates the Chk1p branch). In this case, overexpressing the lnc10883 and Chk1p outcomes a normal cell cycle, precluding the expected arrest by the CHK1 overexpression (**Supplementary Figure 1; Supplementary Table 2**).

## DISCUSSION

### The new cell cycle model robustness

Mathematical modeling can test hypotheses or make predictions of biological data (Noble 2003). Although many *S. cerevisiae* cell cycle models using quantitative and logic mathematical approaches are available (Li et al. 2004; Fauré et al. 2009; Irons 2009; Todd and Helikar 2012; Alcasabas et al. 2013; Rubinstein et al. 2013; Kraikivski et al. 2015), the ones comprise neither all proteins nor lncRNAs and are appropriate to explore only specific cell division mechanisms. Therefore, we developed a new logical model of yeast’s cell cycle, combining the bulk of protein-protein and lncRNA-protein interactions. Afterward, we used transcriptomic data to model the yeast’s cell cycle behavior under ethanol stress.

The cell cycle is a robust system by keeping itself functional even under disturbances (Li et al. 2004; Fauré et al. 2009; Irons 2009; Todd and Helikar 2012; Alcasabas et al. 2013; Rubinstein et al. 2013; Kraikivski et al. 2015). To evaluate the stability of systems’ basin of attraction (a repetitive system’s cycling states) under perturbations is one way to measure the systems’ robustness (Wuensche 2004; Meyers 2020). Simulations indicated that our model is robust since most of the cell cycle mutant phenotypes were adequately simulated, and most random inputs culminated in a cycling behavior. Altogether, we suggest that our model is suitable to test hypotheses concerning the cell cycle.

Although the model had a good performance simulating previous data, the one has two limitations. The DNA must be replicated only once per cycle to maintain cell stability (Mimura et al. 2004); however, our model does not deal with that. Our model correctly predicts mutations related to the FEAR and MEN, essential pathways for proper chromosome segregation in the late anaphase. These pathways rely on Cdc14p release, which may be independent of Cdc15p (Shou and Deshaies 2002). However, the Cdc14p independent release is absent in our model because it decreased the number of correct predictions. Altogether, the model is unfeasible to test hypotheses concerning either DNA replication control or mutations involving the Cdc14p independent release.

### A late cell cycle arrest in LTs induces a better rebound after the ethanol stress relief

Our model and experiments allowed a systemic analysis of the *S. cerevisiae* cell cycle during ethanol stress. The data indicate that LTs have a better population rebound after the severe ethanol stress relief because their cell cycle was arrested closer to the end of cell division (the M phase) compared to HTs (the G1 phase). Therefore, after the stress relief, LTs have a significant time advantage to reestablish and finish the cell division. Indeed, the G1 is the most time demanding phase in the *S. cerevisiae* cell cycle (the whole cycle takes 99, and 142 minutes for mother and daughter cells, respectively) (Brewer et al. 1984). Finally, the population rebound experiment of SEY6210 and BY4742 wild-types corroborated LT outperforms.

Concerning the molecular mechanisms behind the arrests mentioned, our model showed that the SCF complex overexpression in HTs leads to G1 arrest or inviability. Although the literature reports that knockout or conditional expression of SKP1 (express a protein present in the SCF complex) causes arrest in G1/G2 and an inviable phenotype (Connelly and Hieter 1996; Stemmann and Lechner 1996), as far as we know, hitherto here is the first evidence concerning the response of SCF overexpression. We assessed that the lack of Clb1/2p degradation blocks the cell cycle completion, stalling LTs at the M phase. Indeed, the proteolysis of this cyclin, mainly by the APC complex, is crucial for mitosis exit (Pfleger and Kirschner 2000; Wäsch and Cross 2002). Additionally, cells either lacking the APC complex or with a non-degradable Clb2p-form have a high level of Clb2p throughout the cell cycle, turning off the division (Cross 2003).

### The lnc9136 positively work on the cell cycle of SEY6210

The *in silico* overexpression of lnc9136 of SEY6210 outcomes a complete cell cycle. The partial deletion of two different regions of this lncRNA corroborates its positive impact mentioned: one mutation dampened the population rebound after the stress relief, while the other one is inviable.

Concerning the molecular mechanism behind the lnc9136 action, the one indirectly affects the Clb1/2p. The M phase entry relies on the septin formation, which acts at the end of cell division. The conjoint action of Hsl1p and Gin4p, as well as the inactivation of Swe1p, drives the septin formation, which in turn, prompts the M phase entry by a synergic action with the Clb1/2p activation (Barral et al. 1999; Asano et al. 2006). The action of Clb1/2p holds cells at the end of the M phase, then the complete inhibition of this protein is required for the M phase release (Tzeng et al. 2011). Our modeling inhibition of Hsl1P and Gin4p by lnc9136 positively impacts the Swe1p (which inhibits the Clb1/2p) peaking to an M phase release and the cell cycle restart. In this case, the lnc9136 causes a similar effect of HSL1 and GIN4 knockouts, which increases the Swe1p abundance and allows the cell viability (Barral et al. 1999).

LncRNAs can work on cell cycle progression in many ways, including indirect regulation of cyclins, CDKs, and transcriptional factors. For instance, the lncRNA GADD7 expressed in CHO-K1 cells interacts with Tar DNA binding protein 43, leading to the Cdk6p mRNA degradation and preventing the shift from G1 to S (Kitagawa et al. 2013). The role of lncRNAs on cell cycle is deeply studied in human cell lines (including cancers), which can cause cell cycle arrest (sometimes by physical interaction with a diversity of proteins) (Wang et al. 2008; Kotake et al. 2011; Tripathi et al. 2013; Liu et al. 2015; Wei et al. 2016), or even though to promote cell proliferation (Berteaux et al. 2005; Guo et al. 2016; Marín-Béjar et al. 2017).

Despite the scarcity of information about the action of lncRNAs on the yeast cell cycle, some of them are stress-responsive. For instance, under osmotic stress, the Hog1 induces the transcription of the lncRNA Cdc28, driving the increase of CDC28 gene expression and improving cell cycle re-entry after stress relief (Nadal-Ribelles et al. 2012). Experiments showed that four ncRNAs may be associated with ethanol tolerance (Balarezo-Cisneros et al. 2021) and that the severe ethanol stress-responsive lncRNAs work on different pathways such as cell cycle, growth, longevity, cell surveillance, RNA/ribosome biology, among others (Marques et al. 2021).

### The lnc10883 of BY4742 keeps the cell cycle active even under spindle and DNA damages

Our group found that the severe ethanol stress is inducing DNA damage in all strains here studied (unpublished data). However, here we showed that all LTs seem to trigger the cell cycle arrest, while the HTs kept the whole cell cycle even harboring DNA damage.

Two systems synergically control the DNA damage checkpoint, the Rad53p and the Chk1p/Mec1p branches (Chen and Sanchez 2004). However, the activation of a single system is enough to cause arrest to repair the damaged DNA (Melo and Toczyski 2002; Chen and Sanchez 2004; Rock and Amon 2009). Although BY4742 has two DNA damage checkpoint genes overexpressed, the PDS1 (Chk1p branch) and RAD53 genes are down-regulated. Therefore, we claim that the DNA damage checkpoint in this strain is inactive. Interestingly, the lnc10883 inhibits the Mec1p inactivating the Chk1p branch, which also skips the cell cycle arrest. Thus, we suggest that the binding of lnc10883 with Mec1p may be another mechanism to dampen the DNA damage checkpoint.

Additionally, the interaction between lnc10883 with Bub1 inhibits the MCC assembling, a complex essential for inhibiting the APC/C, allowing the cell division even under the putative spindle damage. Indeed, the Bub1p knockout hampers the arrest under treatment with the microtubule polymerization interferent nocodazole, keeping the whole cell cycle even with misaligned kinetochores (Goto et al. 2011).

Previous researches had shown the role of lncRNAs on checkpoints. For instance, the knockdown of the lncRNA PANDA in humans sensitized fibroblasts to apoptosis driven by DNA damage (Hung et al. 2011). DNA damage-induced lncRNAs prompt the DNA damage response in mouse cells (Michelini et al. 2017). The regulation of the DNA damage response mediated by lncRNAs is common in cancers, particularly by modulating ATM, ATR, and p53 signaling pathways (Su et al. 2018).

Here we presented a new model for yeast’s cell cycle and the role of two ethanol responsive lncRNAs in this pathway. Our model allowed analyzing the cell cycle dynamics during the severe ethanol stress. This data, cell culture experiments, and CRISPR-Cas9 mutants evidenced that two lncRNAs probably modulate the cell cycle under the ethanol tolerance and DNA/spindle damage checkpoints by repressive binding to proteins of these pathways. Finally, our model basis is numerous classical genetic data available, the model correctly predicted most mutants phenotypes, and the mutants here generated validated one model prediction. Therefore, we claim that model is suitable for many studies focusing on cell cycle progression.

## ACKNOWLEDGE

We thank the Dr. Arnold Driessen of the University of Groningen, The Netherlands, to kindly donate the plasmids used in this work and the CRISPR-Cas9 know-how knowledge transfer.

## AUTHORS CONTRIBUTION

GTV and LCL conceived the project, performed statistics, and drafted the manuscript. LCL and IRW performed the bioinformatics analysis. APS and LCL worked on wet-lab experiments. All authors agree with the final version of this work.

